# Multi-Fluorescence High-Resolution Episcopic Microscopy (MF-HREM) for Three-Dimensional Imaging of Adult Murine Organs

**DOI:** 10.1101/2020.04.03.023978

**Authors:** Claire Walsh, Natalie A. Holroyd, Eoin Finnerty, Sean G. Ryan, Paul W. Sweeney, Rebecca J. Shipley, Simon Walker-Samuel

**Affiliations:** Centre for Advanced Biomedical Imaging, University College London, London, WC1E 6DD, UK; Physics, Astronomy and Mathematics, University of Hertfordshire, College Lane, Hatfield, AL10 9AB, UK; Cancer Research UK Cambridge Institute, University of Cambridge, Li Ka Shing Centre, Cambridge, CB2 0RE, UK; Department of Mechanical Engineering, University College London, London, WC1E 6BT, UK

**Keywords:** serial-sectioning, HREM, deconvolution, tumor, whole-mount

## Abstract

Three-dimensional microscopy of large biological samples (>0.5 cm^3^) is transforming biological research. Many existing techniques require trade-offs between image resolution, sample size and method complexity. A simple robust instrument with the potential to perform large volume 3D imaging currently exists in the form of the Optical HREM, however the development of the instrument to date is limited to single fluorescent wavelength imaging with non-specific eosin staining. This work presents developments to realize the potential of the HREM to become Multi-fluorescent High Resolution Episcopic Microscopy (MF-HREM).

MF-HREM is a serial-sectioning and block-facing wide-field fluorescence imaging technique, which does not require tissue clearing or optical sectioning. Multiple developments are detailed in sample preparation and image post-processing to enable multiple specific stains in large samples, and show how these enable segmentation and quantification of the data. The application of MF-HREM is demonstrated in a variety of biological contexts: 3D imaging of whole tumor vascular networks and tumor cell invasion in xenograft tumors up to 7.5 mm^3^ at resolutions of 2.75 μm, quantification of glomeruli volume in the adult mouse kidney, and quantification of vascular networks and white matter track orientation in adult mouse brain.

## 1. Introduction

3D optical imaging for large (>0.5cm^3^) intact samples is an increasingly utilized tool in many areas of biomedical research, driven by the desire to understand the 3-dimensional structure of biological systems across multiple biological length scales. All 3D optical microscopy techniques aiming to image large samples must overcome the opacity of tissue to visible light caused by scatter and absorption; tissue clearing and serial-sectioning are two of the most common approaches. Tissue clearing renders tissue optically transparent through de-lipidation and refractive index matching,^[1–3]^ thus enabling visible light to penetrate large (several centimeters) samples fully. Serial-sectioning physically cuts the sample, exposing the deeper tissue layers and removing the need for visible light to penetrate the tissue. Cleared samples can be imaged using techniques such as light-sheet microscopy^[2,4]^ and optical projection tomography (OPT).^[3,5]^ Whilst clearing has been successfully applied to many organs and tissues, the plethora of different protocols can be daunting for new users and a researcher must carefully consider the trade-off associated with each method, e.g. effectiveness of tissue clearing (particularly for large organs) verses stain preservation (endogenous fluorescence and lipophilic dyes), morphological changes, time and complexity.^[6–8]^

In serial-sectioning, samples are embedded in a hard supporting material, such as resin or paraffin, forming a block. Serial sections can then either be cut, mounted, stained and individually imaged, or imaging and sectioning of the block-face can be interleaved (i.e. an image of the block face taken after each successive section is cut). For the former approach, the 3D alignment of the images is a non-trivial challenge due to the significant distortions and misalignments that occur during sectioning and subsequent processing.^[9]^ Conversely, the second approach, serial-section block-face (SSBF) imaging, produces inherently aligned images, thereby overcoming the slice alignment challenge and preventing the potential loss of data through sections damaged during cutting. However SSBF require whole mount staining (similar to light-sheet imaging) and suffer a loss of axial resolution, due to contamination of the image plane by out-of-focus light from below the block’s surface (sub-surface fluorescence).^[10,11]^ The addition of optical-sectioning capabilities such as two-photon and structured illumination to SSBF instruments, has largely overcome the sub-surface fluoresce issue,^[12–15]^ but at the cost of dramatically increasing such instruments complexity and technical specification. These instruments are highly complex, predominantly custom-built and require extensive expertise to align and maintain.^[12,15]^ This puts SSBF imaging beyond the reach for the majority of biomedical researcher labs, and thus creates a niche for a technically simpler SSBF imaging technique for cases where a clearing-based approach is impossible or undesirable. A commercially available SSBF system, the Optical HREM (Indigo Scientific), has been in existence for several years and robust protocols for its use are well established.^[16]^ The system (**Figure 1A**) comprises of a compound microscope head (Olympus MVX10) (as widely used in various light-sheet setups),^[2]^ a light source (LED or arc lamp) with polychromatic mirror and excitation and emission filter wheel for wavelength selection. A CCD camera (users choice) records images after each section. The key feature of the system is the inbuilt automated microtome and software interface for alternate block-face fluorescent imaging and thin section cutting (down to 0.86μm).

**Figure 1.**
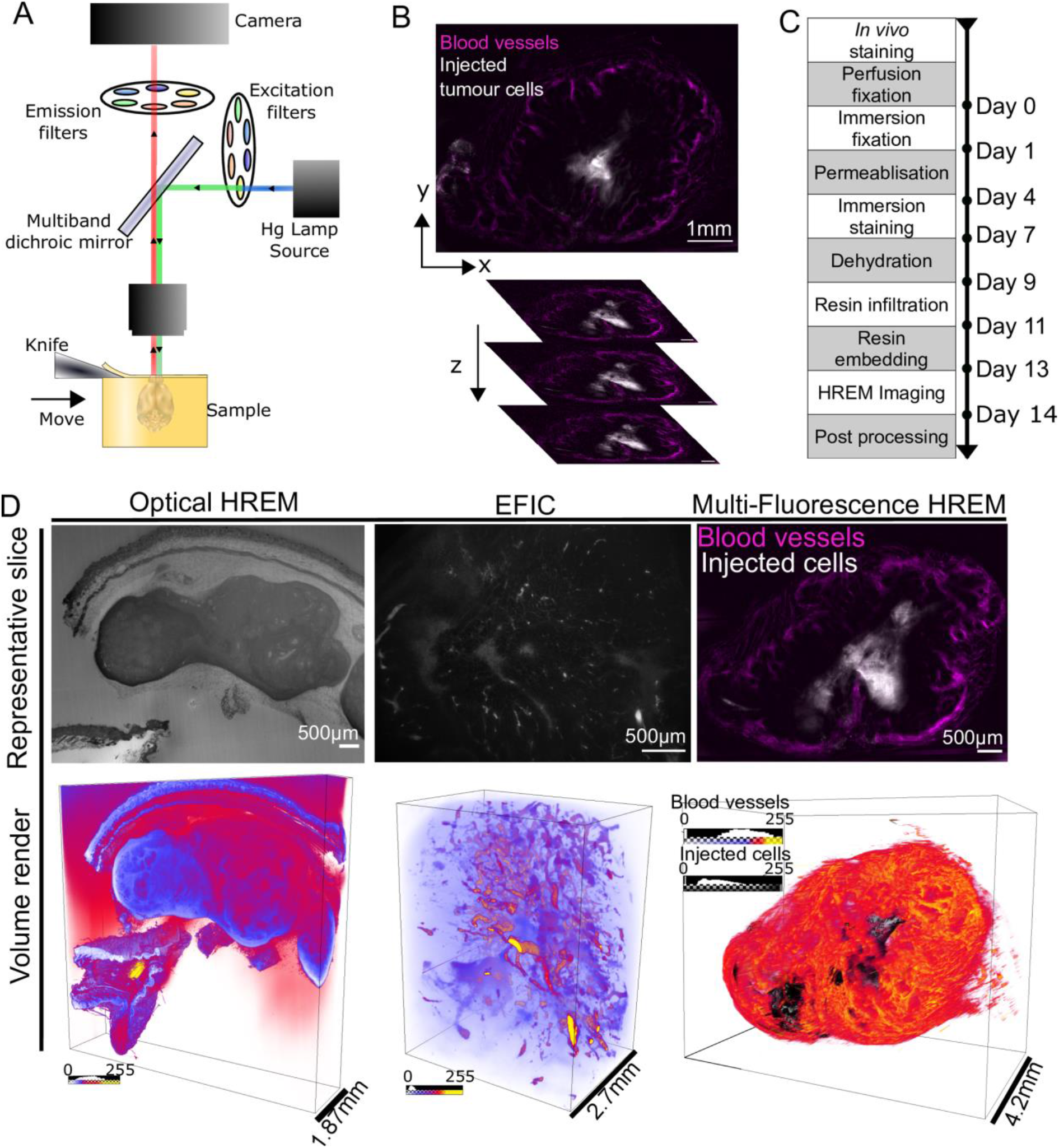
A) The HREM instrument consists of a fluorescent microscope with 1x objective lens (NA:0.25), and a variable zoom which provides fields of view ranging from 25 mm down to 2.3 mm. Biological samples are held within a removable sample holder under the microscope objective on a z-translational stage to enable sections to be cut with a horizontally-aligned, automated sectioning blade. Single-use tungsten carbide blades allow large samples to be cut. The sample is illuminated by a mercury vapor lamp, with separate excitation and emission filters for multiple wavelength imaging. B) Single slices are inherently aligned leading to simple 3D volume rendering of xenograft tumor model. Vascular network as stained by i.v administered lectin and injected cells (stained via the medium-term cell tracing dye CM-DiI) can be clearly identified. C) The MF-HREM sample preparation, acquisition and image post-processing timeline for a typical multi-stained sample. For animal models, the sample is collected following perfusion fixation; in some cases the sample is stained in vivo, prior to fixation. The sample is fixed overnight in PFA before being whole-mount stained (small mount stain or antibody). As almost all candidate resins are immiscible with water, samples must be dehydrated before polymerization. Once staining is complete, the sample is dehydrated using a series of organic solvents, followed by infiltration with a three-part glycol methanlacrylate acrylic (GMA) resin. Finally, the sample is set within the final resin block in the desired orientation and attached to a chuck for mounting to the instrument. Sample imaging with multiple wavelength channels is automated. D) A demonstration of the three HREM imaging methods, utilizing different sources of contrast, in xenograph tumor models. Optical HREM derives contrast from positive, non-specific eosin staining. Tissue autofluorescence provides contrast in EFIC. MF-HREM uses multi-channel fluorescence labelling to visualize multiple specific cellular targets simultaneously.

The Optical HREM instrument and methods were originally developed as a platform for phenotyping mouse models^[17–22]^ and have since been more widely applied.^[23–25]^ The main limitation in its use to date has been the absence of protocols and tools for using multiple targeted contrast agents. The vast majority of HREM to date has utilized a negative contrast eosin approach (via the property of eosin, when bound to eosinophilic proteins, to inhibit the fluorescence of unbound eosin in the embedding resin). This produces images with an appearance similar to the inverse of traditional eosin staining in histology (**Figure 1D**). One exception has been the use of non-fluorescent resin with native autofluorescence.^[26]^ This autofluorescence approach (termed EFIC) was tested on mouse embryos but does not have potential to target specific structures or multiplex stains (Figure 1D). Moreover, the resolution using autofluorescence is far coarser than the eosin staining approach, as no post-processing solutions to recover the axial resolution have been developed.^[26]^

Seeing the potential for this instrument in the 3D imaging field, we have developed the necessary sample preparation techniques and an axial resolution post-processing method to enable dual fluorescent labelling, multi-channel imaging and quantification of specific biological structures in tissue samples >0.5um^3^, at resolution up to 2.75 µm using the Optical HREM instrument (**Figure 1B**). In this paper, we present the optimization of the MF-HREM methodology, including stain penetration and resin embedding. We also describe a two-stage approach to recovering axial resolution, firstly using an opacifying agent (Orasol Black (OB)) to limit light transmission into the sample, and secondly using deconvolution in post-processing with a point-spread-function (PSF) estimated from the image stack itself. Finally, we demonstrate the wide applicability of MF-HREM by: 1) quantifying glomeruli volume in adult mouse kidneys; 2) segmenting vascular networks and invasive cells in a mouse tumor xenograft model; and 3) segmenting vascular networks and quantifying white matter tract orientation in a mouse brain. We show here that these developments greatly broaden the potential applications of HREM and provide a large-volume 3D imaging platform that is accessible to a wide range of researchers.

## 2. Results

### 2.1 Optimization of sample preparation

The pipeline for MF-HREM is straightforward and does not require specialist equipment (Figure 1C). Each stage of the pipeline requires optimization for a specific experiment (organ and stain combination) and we have performed this optimizations for a variety of adult mouse organs and experimental conditions. These optimized protocols can serve as a starting point for other experimental conditions and demonstrate the breadth of potential applications for MF-HREM.

#### 2.1.1 Staining

As MF-HREM requires whole mount staining prior to dehydration and resin infiltration, stain compatibility with the process must be established. We have tested a wide variety of stains and assessed their compatibility with various dehydrants and resins. We have particularly focused on providing robust counter-stains for cell nucleus (HCS nuclear Mask), cytoplasm (HCS cell mask) and cells membrane (Wheat Germ Agglutinin), as well as for vascular staining lectin-dyelight conjugate (a full list of stains can be found in **Table S2**). It is noteworthy that the lipophilic stain, CMDiI, is compatible with MF-HREM where it is not with many clearing techniques.^[7]^

Whole-mount staining requires homogenous and rapid stain penetration, which can be improved by increasing tissue permeability. Four methods to increase the permeability of tissue samples that have been previously used^[1,5,37,38]^ were compared: freeze-thaw, proteinase K (P[K]) digestion, iDISCO (which combines several mild detergents)^[1]^ and saponin. The comparison of the four methods on adult mouse kidneys showed that saponin-treatment significantly increased stain penetration, compared with the control case (p=0.04) (paired t-test N=4). The iDISCO method also increased stain homogeneity (p=0.055) compared to control kidneys **(Figure 2A-C)**.

**Figure 2.**
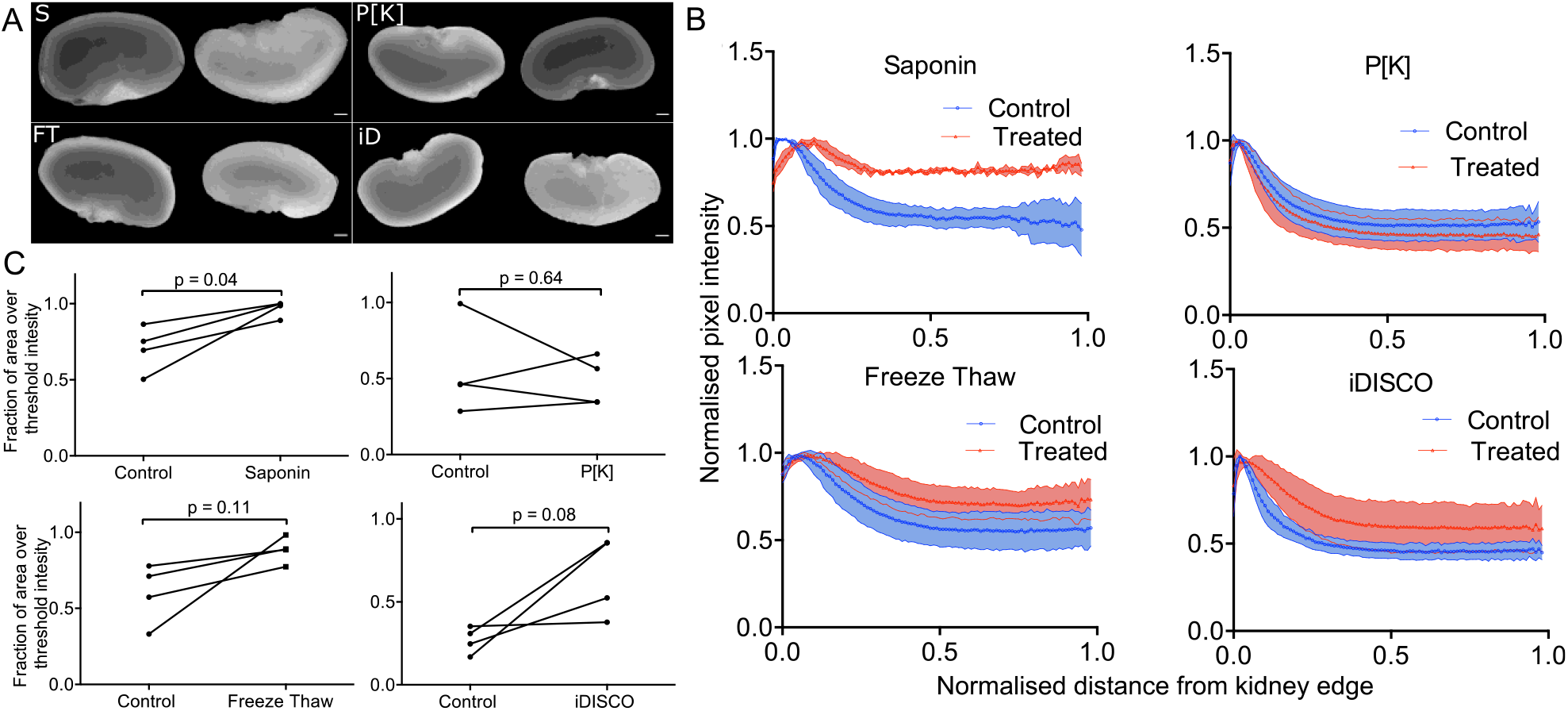
Optimization of stain penetration in adult mouse kidney samples. Four methods for improving stain penetration are compared: saponin treatment, proteinase [K] digestion, freeze**-**thaw and iDISCO. For each method, four treated and four control kidneys were investigated, with one kidney from each animal used as a control for the contralateral kidney. A) Representative images of the kidneys, imaged as described in section 5.2, alongside the control (contralateral kidney on the left and the treated kidney on the right) (scale bar, 1 mm). B) shows the normalized MF-HREM signal intensity profile as a function of radial distance from the kidney edge. C) shows the fractional area of the kidney section image, above a threshold value (the same threshold was used for each treated kidney and matched control). Results of paired t-test analysis demonstrates that saponin treatment significantly increased stain penetration (p<0.05).

Alternative staining routes, such as intra-vascular (i.v.) injection are also compatible with MF-HREM in animal models. For vascular staining, use of i.v. injection of fluorescently-conjugated lectins is effective with the MF-HREM pipeline across a range of organs as shown in **Figure 5, 6 and 7**.

**Figure 3.**
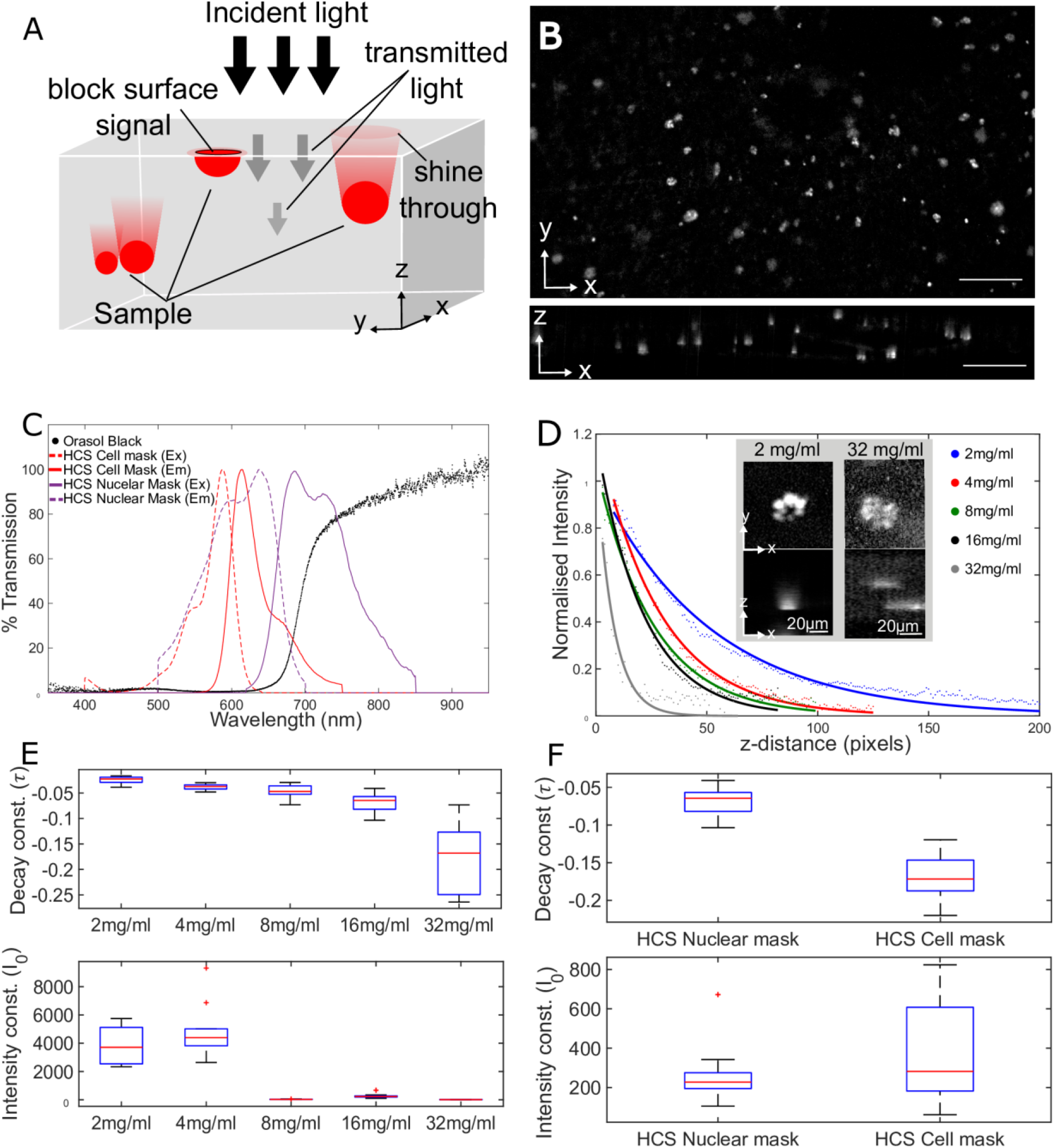
Characterization of OB as an opacifying agent to reduce sub-surface fluorescence. A) Diagram demonstrating the origin of sub-surface fluorescence. B) A representative image of cells in an in vitro 3D culture stained with HCS Nuclear Mask with 16mg/ml of OB. The comet tail artefact can be seen in the xz plane image (scale bar is 100 µm). C) Graph showing the measured transmission spectrum of OB (0.1 mg/mL, 4 mm path length), as well as two tested commercial stains HCS Nuclear Mask and HCS Cell Mask (spectrum from manufacturer). It can be seen that OB has low transmission in the 400-625 nm range which rises steeply in the 625-700 nm range. The HCS Cell mask spectrum falls almost entirely within the low transmission band of OB whereas the emission of HCS Nuclear mask falls in the section of steep increase in transmission. D) A single exponential fit to the mean intensity profiles for 10 xz plane ROIs taken of single cells in 3D *in vitro* cell culture stained with HCS Nuclear Mask of increasing OB concentration (2, 4, 8, 16, 32 mg/ml) R^2^ values (0.976, 0.994, 0.989, 0.982, 0.803) respectively. Pixels correspond to a distance of 1.72 µm (slice thickness). The inset shows representative single cells with the minimum and maximum OB concentration. E) Values of exponential decay constant (*τ*) and initial intensity (*I*_*0*_) are shown for each of the concentrations (see section 5.5) and show the expected decrease in decay constant with increasing OB concentration. There is also a large decrease in intensity at 8 mg/ml and higher OB concentrations. F) The same fitting parameters are compared in HCS Cell Mask stained sample and HCS Nuclear mask stained sample at 16 mg/ml OB. As expected from the transmission spectrum, the decay constant is more negative in the HCS Cell Mask case than the HCS Nuclear Mask case.

**Figure 4.**
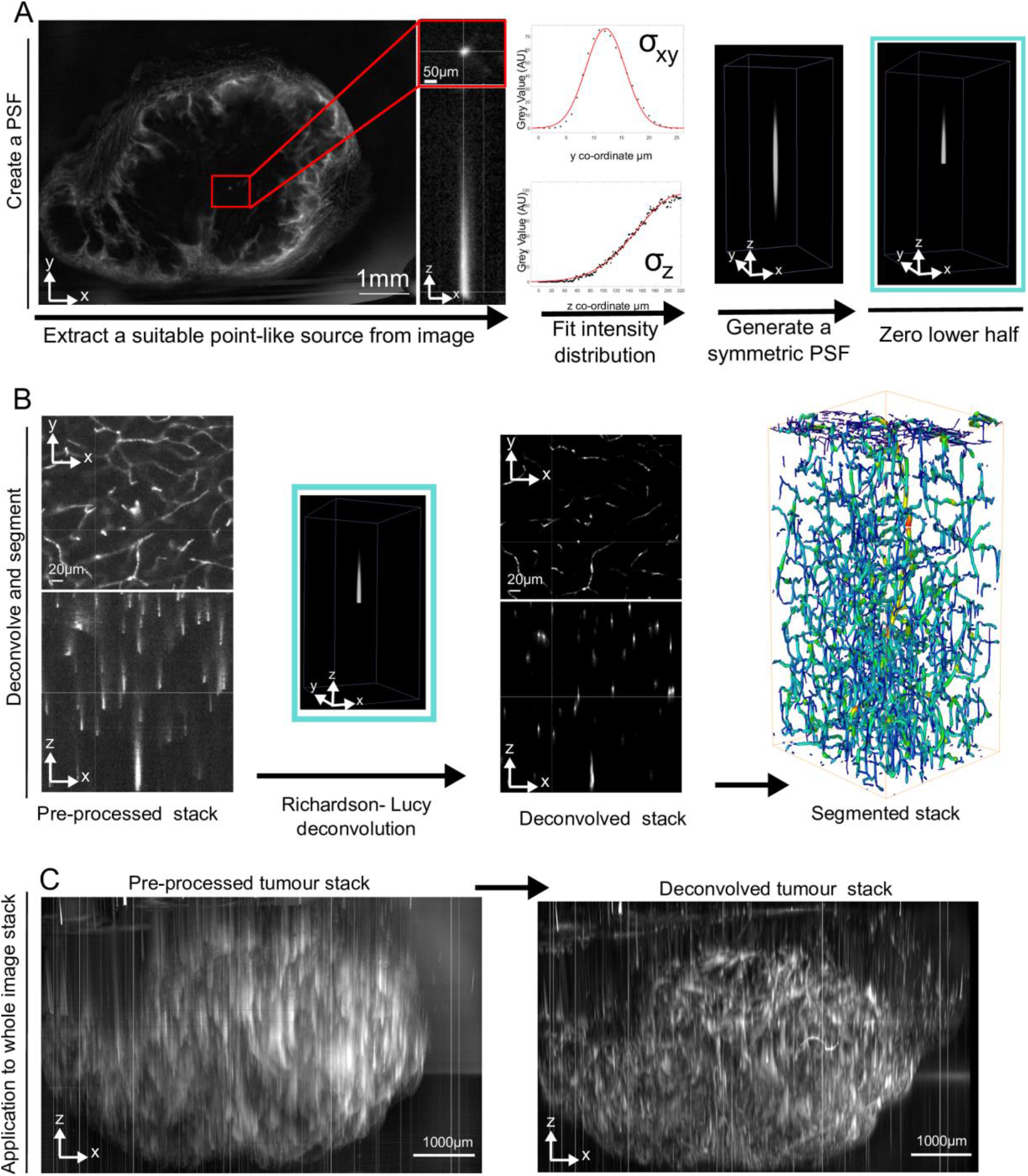
The Image processing pipeline. A) Showing the pipeline for the extraction of a PSF from an image stack of a subcutaneous tumor with microvascular stained via injection of Lectin-Dyelight649. A suitable point-like source is found in the image stack, this is cropped from the image and a Gibson & Lanni PSF model is fitted to the data by minimizing the difference between the measured and synthetic FWHM. The synthetic (symmetric) PSF is generated with PSFGenerator.^[45]^ This PSF is then half zeroed to create the final PSF. B) The deconvolution of the pre-processed image stack using the PSF. A Richardson-Lucy method^[29]^ is used to deconvolve the stack creating images that can then be segmented and quantified using various methods dependent on the biological context. C) Application of the PSF estimation and deconvolution method to a whole tumor vascular network improves axial resolution.

**Figure 5.**
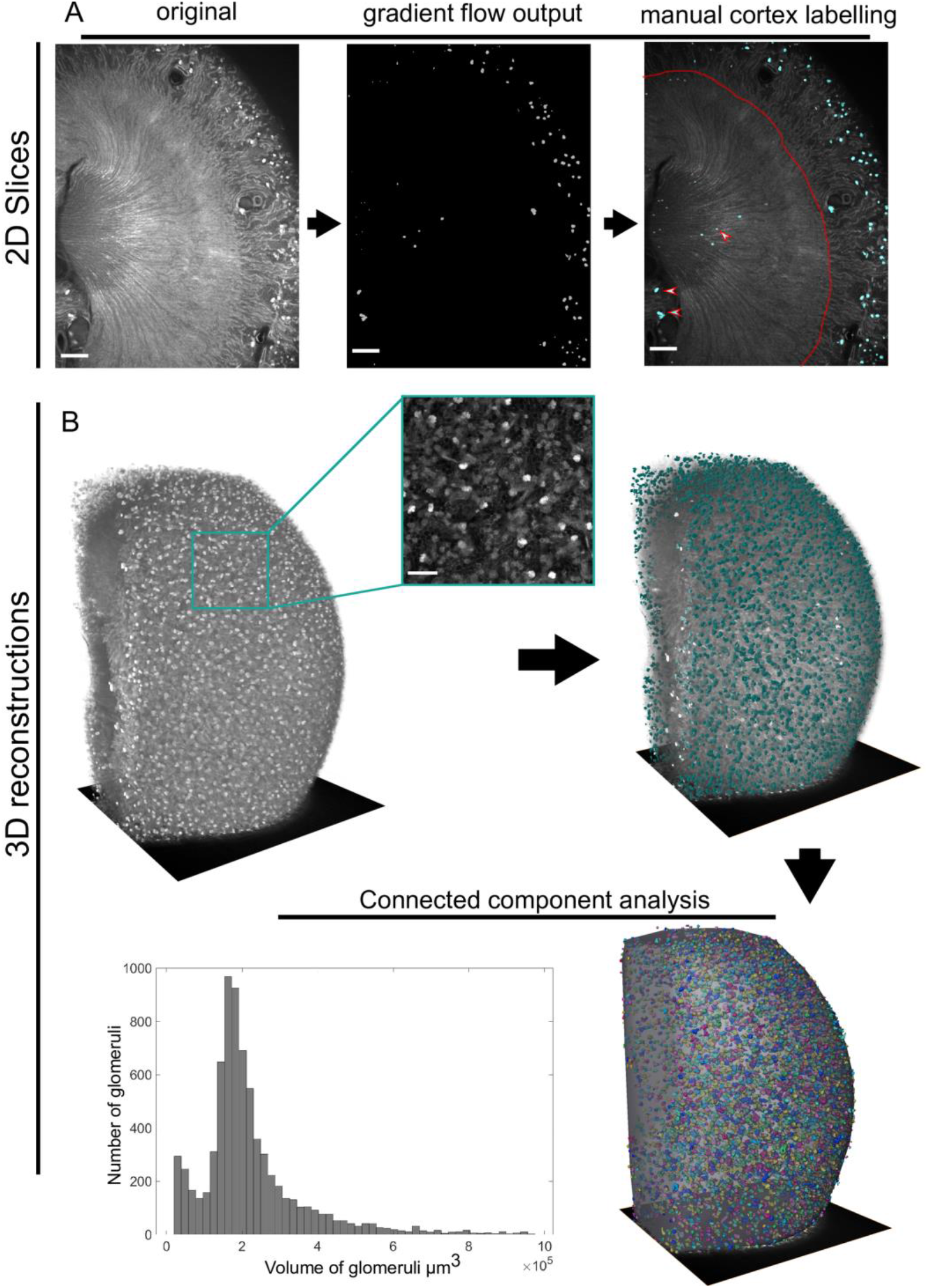
Quantification of glomeruli number from the Lectin-Dyelight649 channel of a multi-stained murine kidney. A) From left to right, 2D slices showing: the unprocessed image, the segmented image produced by gradient vector flow algorithm, and finally, the manually-corrected image after removal of structures that were segmented (white/red arrows) but fall outside the kidney cortex (red line). Scale bars 500 µm for all 2D slices. B) 3D view of the image processing. The original image stack with inset showing the detail which can be seen on the kidney surface (scale bar 200 µm), then the segmentation via gradient flow algorithm in Vaa3D^[30–32]^ and manual exclusion of points not in the cortex. The final step is the outcome of the connected components analysis and hence quantification of glomeruli number, and volume distribution. These data show the expected distribution and size of glomeruli for heathy wt mouse as compared with other techniques.^[47,48]^

**Figure 6.**
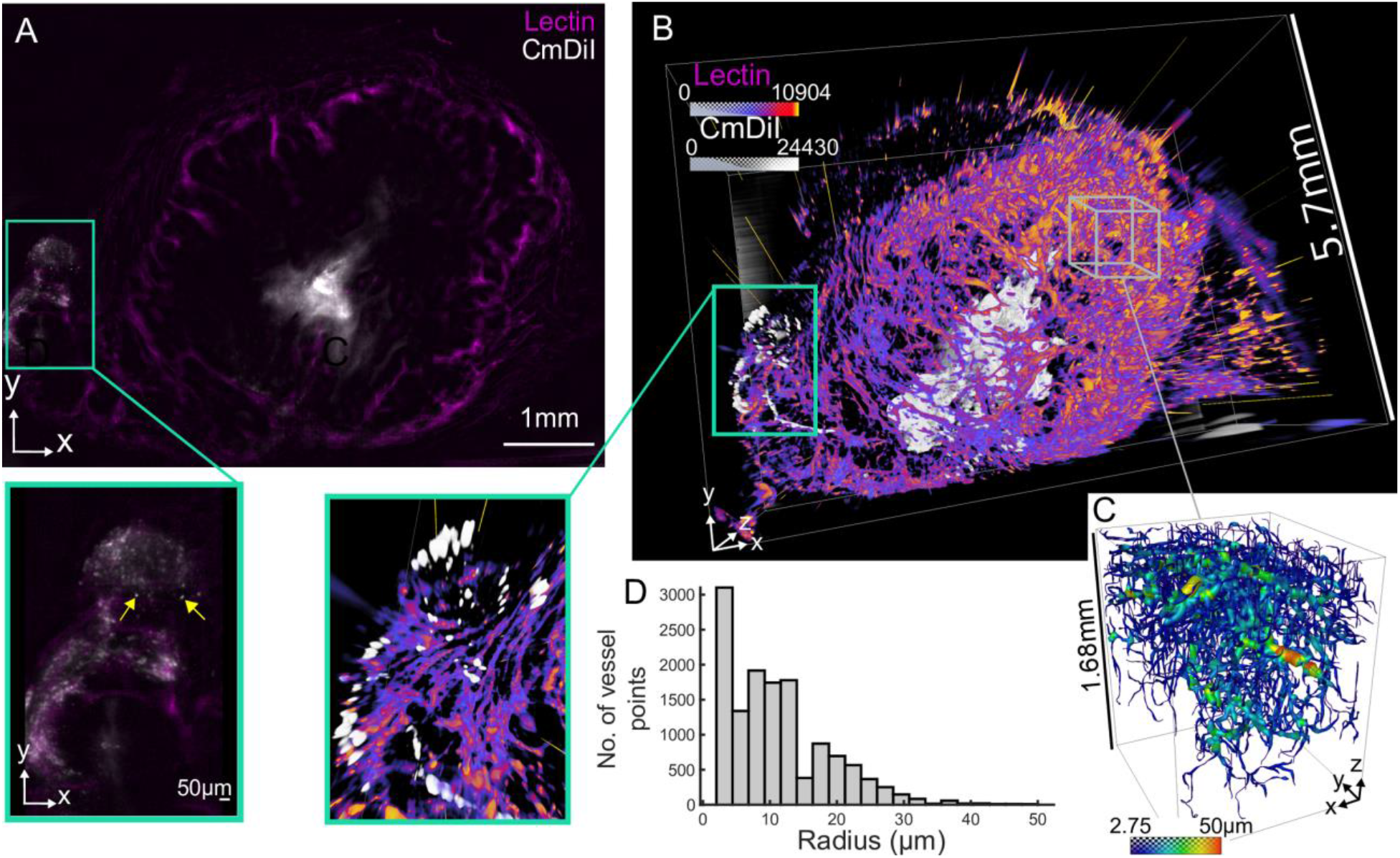
Xenograft tumor model analysis. A) Shows a single representative slice with the two stains: CMDiI for injected cell tracking (white), and Lectin-Dyelight 649 conjugate for microvascular staining (magenta). The inset shows a digitally zoomed in portion of the image where small, approximately circular structures of cell size are indicated by yellow arrows. B) A 3D rendering of the two channels shows the tumor in its totality and the spatial arrangement of the injected cells within the vascular network. The highly perfused rim can be clearly seen. The inset shows the same digitally zoomed section as the 2D slice inset. C) Segmentation of a section of the vascular network following the PSF extraction described earlier. In this case the APP2 algorithm from the Vaa3D neuron tracing plugin set was used to segment and skeletonize the deconvolved image stack.^[30–33]^ D) Shows the histogram of the vessel radii from C.

**Figure 7.**
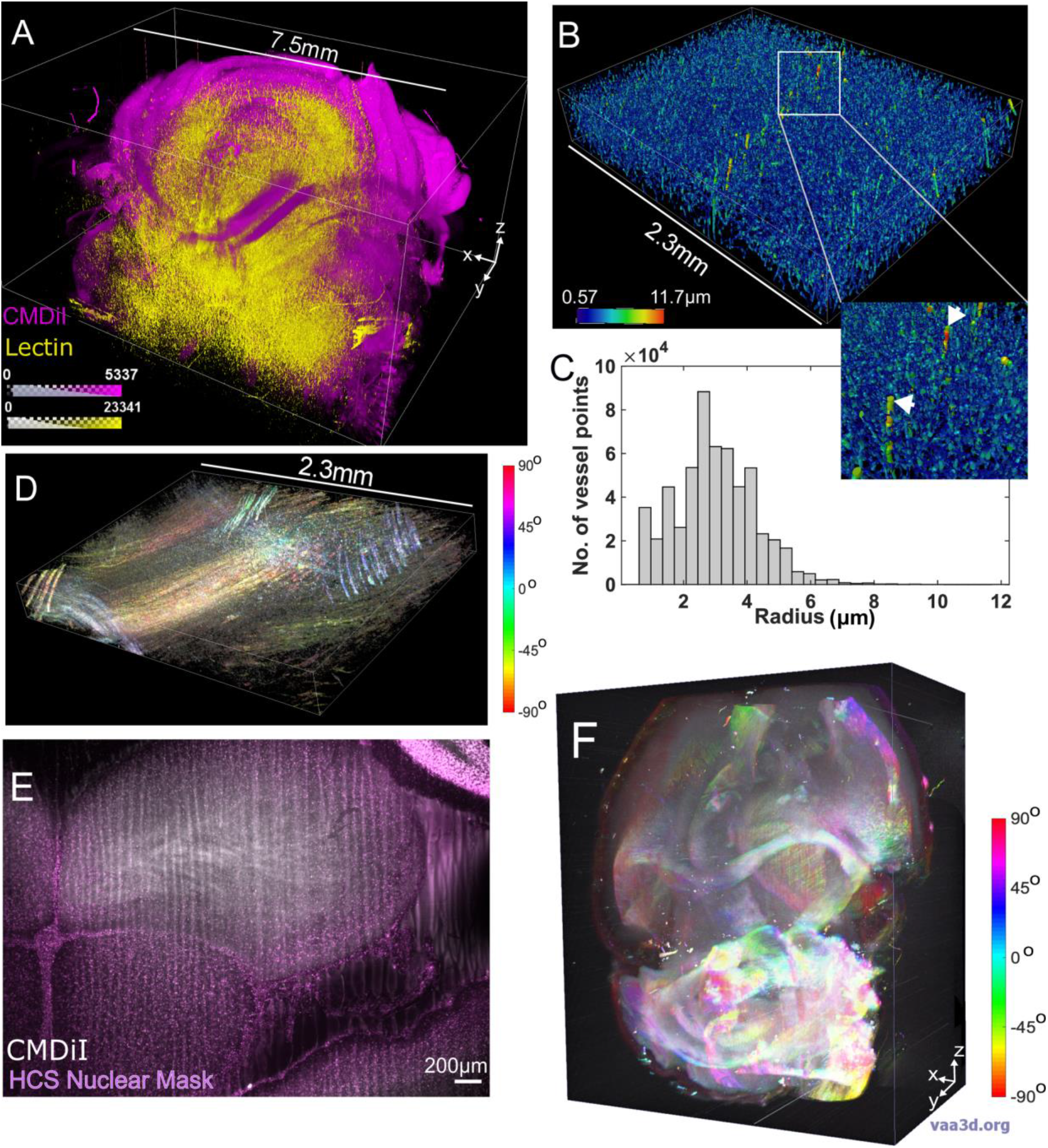
MF-HREM of two brain samples, one stained with CMDiI Ex/Em 553/570nm and Lectin-Dyelight 649 (Ex/Em 649/700) and a second stained with CMDiI and HCS Nuclear Mask (Ex/Em 638/686). A) The volume overlay of both the CMDiI (Magenta) and the Lectin (yellow). Color bars show pixel intensity. B) A high resolution sub-volume of the vasculature channel successfully segmented and skeletonized using the MF-HREM processing pipeline and the MOST tracing algorithm implemented in Vaa3D. C) The histogram of radii distributions. D) The same sub-volume in the CMDiI/ white matter channel; here an orientation analysis has been performed via calculation of the structure tensor of the image using a Gaussian gradient and 8 pixel window size. The hue, saturation and brightness denote the orientation, coherence and original image brightness respectively. The color bar shows the angle represented by the hue. E) A polar histogram of D where all orientations with a coherence greater than a threshold value of 0.2 are displaced. It can be seen that there appears to be an even distribution of local orientations over this sub volume. F) Volume overlay of the CMDiI channel (white) and the HCS nuclear mask channel (magenta) for a different brain sample. G) A single slice imaged at higher resolution in which nuclei can be clearly seen. H) Orientation of white matter mapped over the whole brain. It was computed as in D, but using a smaller window size of 4pixels. The orientation can be seen to clearly follow the expected white-matter tracts.

#### 2.1.2 Resin infiltration and embedding

After staining, samples must be dehydrated and embedded in resin to provide mechanical stability during sectioning. Various commercial resins are used in histology, however, as these resins are designed to be manually cut and subsequently stained, they are not optimized for automated, thin sectioning, fluorescence preservation or, in many cases, large samples.^[39]^ We investigated the compatibility of five commercial resins, which covered the three broad chemical categories for hard resins: methacrylate resins (Technovit 7100, Technovit 8100 and Lowicryl HM20), epoxy resin (Spurr) and acrylic resins (LR White).^[39–41]^ Resins were assessed for the time taken to set, their compatibility with an opacifying agent OB (see section 2.2.1 for further details), for their cut quality and for hardness.

Technovit 8100, provided the best combination of cut quality, hardness, time to set and compatibility with OB (**Figure S1** and **Table S3**). The addition of OB to Technovit 8100 showed that resin hardness decreased with increasing concentrations of OB, but that this could be counteracted by increasing the concentration of the resin’s secondary catalyst (Figure S1).

Lastly, infiltration/embedding times for a variety of adult murine tissues were optimized to provide protocols for different sized organs (**Table 1**).

**Table 1.** Optimization of dehydration and embedding times for samples or various sizes.

| Reagent | >60 mm<sup>3</sup><br>Embryo (up to<br>E12.5) Organoid or<br>3D culture | 60-400 mm<sup>3</sup> e.g.<br>sciatic nerve or<br>embryo E14.5 to P1 | 0.4-1 cm<sup>3</sup><br>e.g. heart, kidney,<br>brain, tumor | >1 cm<sup>3</sup> |
| --- | --- | --- | --- | --- |
| Acetone 50% | 1hr | 6hrs | 12hrs | 24hrs (refresh at<br>12hrs) |
| Acetone 70% | 1hr | 6hrs | 24hrs<br>(refresh at 12hrs) | 24hrs (refresh at<br>12hrs) |
| Acetone 80% | 1hr | 2hrs | 2hrs | 12hrs (refresh at<br>6hrs) |
| Acetone 100% | 15mins x3 | 1hr x3 | 2hrs x3 | 3hrs x3 |
| 50:50 Acetone: infiltration sol | 2hrs | 12hrs | 12hrs | 24hrs |
| 25:75 Acetone: infiltration sol. | NA | 12hrs | 12hrs | 24hrs |
| 100% Infiltration sol. + vacuum | 2hrs<br>include OB | 12hrs<br>include OB | 24hrs (refresh at<br>12hrs) include OB<br>after refresh | 48hrs (refresh at<br>24hrs) include OB<br>at refresh |

### 2.2 Minimizing subsurface fluorescence

#### 2.2.1 Minimizing sub-surface fluorescence with Orasol Black

In any widefield microscopy technique fluorophores that are above or below the focal plane may be excited and their emission captured as out-of-focus light at the focal plane (**Figure 3A**). For MF-HREM and other SSBF techniques, this effect is asymmetric due to the physical sectioning of the sample above the focal plane. This leads to a characteristic comet-tail like artefact in the axial plane which prevents image segmentation, as seen in **Figure 3B**. To reduce this so-called sub-surface fluorescence, we investigated the addition of an opacifying agent (OB) to the embedding resin. The aim was to reduce the penetration of incident light into the block and absorb emitted photons from beneath the block’s surface thus reducing subsurface fluorescence.^[42]^

The extent to which OB reduces subsurface fluorescence is wavelength-dependent. The transmission spectrum of OB is shown in **Figure 3C**. It has a broad absorption in the 450nm to 650nm range, with a steep increase in transmission in the near infrared range (>700nm). We measured the decrease in subsurface fluorescence with increasing OB concentration at wavelengths of 705 nm and 600 nm. With increasing concentration of OB, we found a decrease in decay constant *τ* and a visible reduction in comet-tail artefact (**Figure 3E**). The initial intensity *I*_*0*_ shows a sharp decrease at 8 mg/ml, which can be seen in the image stacks as a reduction signal-to-noise ratio, and the highest OB concentration (32 mg/ml) provided the greatest decrease in subsurface fluorescence, but, as previously discussed, this affected resin polymerization. A balance between minimizing subsurface fluorescence, achieving adequate signal-to-noise for image processing and optimum resin setting must be found. Concentrations between 1-4 mg/ml were found to be optimum depending on the organ type structures imaged and fluorophores used (see section 3 and **Table S1**). As expected, transmission spectra showed that sub-surface fluorescence was lower for HCS Cell Mask (Em 600 nm) than for HCS Nuclear Mask (Em 705 nm), for the same concentration of OB (**Figure 3F**).

#### 2.2.2 Minimization of shine-through artefact with image post processing

As OB cannot be used to entirely eliminate subsurface fluorescence, we investigated post-processing strategies to deconvolve the collected signal and enable segmentation. Deconvolution requires estimation of the system PSF, which can be achieve by direct measurement of sub-resolution fluorescent beads, synthetically generated from known system parameters or estimated via blind deconvolution methods.^[43]^

In widefield or confocal microscopy, out-of-focus light contamination comes from both above and below the focal plane resulting in a symmetric PSF. In SSBF, the PSF is highly asymmetric and hence it cannot be well estimated by blind deconvolution techniques or by a range of widely used optical models.^[10,11,44]^ Experimentally measuring the PSF is time-consuming (PSFs must be obtained for each new sample and each imaging wavelength) and require a high signal to-noise ratio.^[43]^ Due to these practical difficulties, we sought a method that uses structures from within the image stack estimate the PSF.

It has previously been shown that small structures in two-photon data could be used in place of sub-resolution beads to estimate a PSF.^[46]^ Using this approach on MF-HREM data was unsuccessful due to the poorer signal-to-noise ratio of MF-HREM compared to two-photon microscopy. However, measurements of the full width half maximum (FWHM) for small structures were used to parameterize a synthetic PSF model (**Figure 4A**).

A synthetic PSF was created using the Gibson and Lanni Model, this model was fitted to the mean measured FWHM of several selected small structures (Figure 4A) (see Methods for fitting details). The lower half of the synthetic PSF was zeroed to replicate the asymmetry caused by serial-sectioning. This approach provides the practical advantages of using the image stack for PSF estimation, with the benefit of synthetic signal-to-noise ratio. The asymmetric PSF was then incorporated into the widely used Richardson-Lucy deconvolution algorithm, an iterative method implemented in many open source software repositories^[10,29]^ (**Figure 4B**). Our approach was applied to image stacks of different magnifications, wavelengths and concentrations of OB (see **Table S1** for imaging parameter values). In all cases there is improvement in axial resolution and in the case of higher OB concentration and high magnification the approach successfully enables segmentation of large vascular networks (section 3.2). For lower concentration of OB and lower magnification (**Figure 4C**) our approach enables automated quantification of large subsections of the image (section 3.2) and improves the comet tails artefact appearance (Figure 4C).

## 3. Applications

### 3.1 Glomeruli number and volume in adult mouse kidney

The nephron is the microscopic structural unit of the kidney, comprising of two parts: the renal tubule and renal corpuscle. The renal corpuscle contains a cluster of capillaries known as a glomerulus. Nephron number has been used in both human and animal models as a biomarker of both renal and cardiovascular disease,^[47]^ however total nephron number and glomerulus size are challenging to estimate from 2D sectioning techniques. **Figure 5** shows an adult mouse kidney stained with a vascular marker, prepared and imaged using our MF-HREM pipeline. Glomeruli were segmented in the final image stack. MF-HREM pixel size was 2.17 µm lateral, or ‘in-plane’, and 2.58 µm axial (voxel volume 12.14 µm^3^), enabling identification of glomeruli which were found to have a minimum volume of 2.4×10^4^ µm^3^.

Our MF-HREM analysis measured 100 glomeruli per cubic micrometer of kidney volume, with median glomeruli size of 1.88 ×10^5^ µm^3^ and mean of 2.2×10^5^ µm^3^. The number of glomeruli per unit kidney volume is consistent with distributions for wild type adult mice performed with light-sheet^[48]^ (100-140 glomeruli per unit volume of kidney), and also MRI^[47]^ (74-90 glomeruli per unit volume of kidney). Our median size is also within the bounds of those measured by lighsheet,^[48]^ stereology and MRI^[47]^. This demonstrates the ability of MF-HREM to provide accurate quantitative information on organotypic functional units in adult mouse organs.

### 3.2 Imaging tumor blood vessels and cell invasion with MF-HREM

Widely reported hallmarks of cancer include deregulation of angiogenesis and active invasion/metastasis.^[49]^ Angiogenic disruption is evident in solid tumors through highly complex and disordered blood vessel networks. These networks are responsible for the distribution of nutrients and drugs,^[3]^ and are involved in bidirectional signaling that can stimulate cellular invasion and, ultimately, lead to metastasis.^[50]^ Multiplexed 3D imaging of whole blood vessel networks with tumor cell invasion can be used to interrogate the spatial relationship between deregulated angiogenesis and cell invasion.

**Figure 6** shows MF-HREM images from a subcutaneous xenograft tumor mouse model initiated from the FaDu hypopharyngeal cancer cell line, where both tumor cells and the blood vessel network have been fluorescently labelled. Both stains are clearly visible throughout the image stack (**Figure 6A**), and 3D reconstruction of the entire data set for both channels reveals the dense, branching vasculature at the periphery of the tumor, and the labelled cells primarily in the tumor center, which appeared to be non-perfused due to an absence of vascular signal (**Figure 6B**). Single cells or cell clusters are also visible (yellow arrows in Figure 6A inset) in a section of tumor separate from the main bulk. Whilst it is unclear whether labelled cells are viable (which would require a different reporter strategy) these results demonstrate the ability of MF-HREM to quantify the 3D location of injected cells in tissue volumes ∼1cm^3^ several weeks after injection.

For the vascular imaging stain, **Figure 6C** and **6D** show a sub-section of the tumor vasculature that has been segmented following image processing with our deconvolution approach. The chaotic nature of the vasculature is clear from both the appearance and from the wide range of vessel sizes (Figure 6D). Such segmented vascular networks can be used in simulations of drug delivery^[3]^ and for investigating tumor vessel growth mechanism.^[51]^

### 3.3 Imaging brain microvasculature and white matter tracts in a mouse brain with MF-HREM

Healthy brain development and function is dependent upon the intricate networks and of both cerebrovascular and white matter fibers. Changes in the structures of these networks are key indicators for a number of degenerative neurological diseases including Alzheimer’s and Parkinson’s.^[4]^

Visualizing these structures in 3D with MF-HREM could be used to improve our understanding of disease, or potentially validate *in vivo* imaging tools such as MRI.

**Figure 7** shows the application of MF-HREM in two instances: i) where a brain is dual labelled with CMDiI as a white matter-marker and lectin-Dyelight649 as a microvascular stain (**Figure 7A-D**); ii) where a brain is dual labelled with CMDiI as a white-matter marker and HCS Nuclear Mask as a marker for cell distribution (**Figure 7E-7F**). Figure 7B shows segmented vasculature from a high-resolution sub-volume of the medulla. The observed microvasculature has a mean radius of 3.01±1.25 µm, with a large number of small vessels (1.6-12 µm diameter) oriented parallel the brain surface and a smaller number of larger (12-22 µm diameter) descending arterioles perpendicular the brain surface (white arrows in Figure 7B inset). This distribution is expected anatomically and accords well with those measured by other serial-sectioning modalities, e.g. MOST (2.8-2.4 µm for matched sub-regions),^[52]^ and for clearing techniques.^[53]^ It should be noted that no large vessels (>22 µm) are visible in MF-HREM data, likely due to the preferential binding of lectin to microvasculature over larger vessels as noted previously.^[54]^

An additional feature of interest in brain microstructure is white matter tract orientation. Previously, CMDiI has been used on histological sections of mouse brain to validate tractography from DW-MR in 2D.^[55]^ CMDiI effectively stains white matter tracts due to its lipophilic nature and can be imaged with MF-HREM allowing orientation analysis on the 3D volumes. Figures 7D and 7F show white matter orientation analysis for a small brain section and whole mouse brain respectively. Orientation is quantified by the structure tensor of the image which was estimated using the ImageJ plugin OrientationJ.^[34–36]^ The color bar shows the angle represented by the hue.

Cell distribution and density are also important marker of neurological development and pathology. Figure 7E shows a single slice at a higher magnification from the imaging volume in Figure 7F, cell nuclei can be distinguished and demonstrate that nuclear staining has successfully been retained throughout processing. Further segmentation requires montaging of higher resolution images.

These data show that MF-HREM can be used in adult mouse brain as a tool for visualizing vascular networks, white matter tracts and cell distribution. Quantitative vessel analysis can be performed on higher resolution data, and orientation analysis on whole brain images.

## 4. Conclusion

In this work, we have shown the development of sample preparation and image post-processing in MF-HREM. We have demonstrated its applicability in adult mouse organs through a range of staining and quantification approaches.

Reducing the impact of sub-surface fluorescence has been a key factor in realizing the potential of MF-HREM. Our current solution has combined two approaches: physically limit light transmission through the sample, and deconvolution in post processing. Our deconvolution approach uses a sample specific PSF, whilst circumventing the problem of poor signal-to-noise that would occur if the PSF is used directly from the image stack. Likewise, our optimization of tissue-specific staining and embedding protocols provides a solid foundation from which researchers hoping to apply this technique can build.

Further improvements to MF-HREM are possible in several areas of the imaging pipeline. The resin embedding and staining protocol can be improved by optimization to a specific biological problem. For example, a custom mould can be used for repeated preparation of similar samples. Additionally, whilst dehydration and embedding times cannot be easily decreased without compromising the final imaging, automation of the process using an automated histological sample processor could improve consistency and enable faster protocol optimization for a specific application.

Diffusion staining alone will always struggle to achieve homogenous stain penetration in a timely manner. Even with increased stain penetration via saponin treatment, inhomogeneous staining is evident. This challenge is common to all whole-mount imaging techniques, thus significant research effort continues to be directed at overcoming this, e.g. the use of nanobodies, genetically encoded reported and perfusion staining.^[56,57,58]^

The two interdependent approaches used here to circumvent sub-surface fluorescence could undergo more development to widen MF-HREM application. To further reduce light transmission, alternative dyes could be incorporated into the resin or shorter wavelength incident light used.^[59]^ Further improvements to the deconvolution could be made through more complex PSF models, for example Gaussian mixture models^[60]^ or spatially variant PSFs.^[61]^

Both optical clearing and serial-sectioning come with their own specific challenges. Approaches which keep the sample intact (thus requiring optical clearing), are limited in sample size and resolution by the working distance of the microscope objective lens^[2]^ and the need for broader light sheets to penetrate greater tissue depth.^[12]^ For applications where a serial-sectioning approach would be better suited, uptake of serial-sectioning techniques has been limited by the technicality of the custom-built instruments,^[15,62]^ leaving a niche for a robust, commercially available serial-sectioning technique.

In this work, we have developed a pipeline for performing MF-HREM, which enables 3D, multiplexed fluorescence imaging of large tissue samples (> 0.5 cm^3^), at high resolution. MF-HREM is a block-facing technique, using a commercially available system (Optical HREM, Indigo Scientific, UK), that overcomes sub-surface fluorescence using a combination of an opacifying agent and image deconvolution. This technique could find wide application through its avoidance of optical sectioning, tissue clearing or the need for a custom-built instrument.

## 5. Methods

### 5.1 Animal models and perfusion fixation

All animal studies were licensed under the UK Home Office regulations and the Guidance for the Operation of Animals (Scientific Procedures) Act 1986 (Home Office, London, United Kingdom) and United Kingdom Co-ordinating Committee on Cancer Research Guidelines for the Welfare and Use of Animals in Cancer Research.^[27]^

For perfusion fixation, animals were euthanized via intraperitoneal (i.p.) injection of 100 mg kg^−1^ sodium pentobarbital (Animalcare, Pentoject) diluted in 0.1 ml phosphate buffered saline (PBS). Once anesthesia was confirmed, surgical procedures for cardiac perfusion were performed for systemic clearance of blood. Heparinized saline (20ml) with 1,000 IU ml^−1^, maintained at 37 °C) was administered with a perfusion pump (Watson Marlow, 5058) at a flow rate of 3 ml min^-1^ to mimic normal blood flow. After the complete drainage of blood, mice were perfused with 20 ml of 4% paraformaldehyde (PFA, VWR chemicals 4°C). Organs were then removed and fixed for 12-24h in 4% PFA at 4 °C.

### 5.2 Stain penetration

A total of 16 mice (N=4) between 10-23 weeks old of various strains were used. Mice were perfuse fixed as described above and both kidneys were removed. Each animal was randomly assigned to one of the four groups (saponin, freeze-thaw, iDISCO and proteinase [K] digestion) and for each animal one kidney was randomly assigned to treatment and the contralateral kidney retained as a matched control. In the saponin, iDISCO and proteinase [K] groups, the control kidney was maintained in PBS and at the same temperature as the treated kidney. For the freeze-thaw group, the control kidney was dehydrated and rehydrated through the same methanol series but with no freeze-thaw cycles applied. Details of the timing and solutions composition for each of the four groups is given in Supplementary Methods.

After treatment, all kidneys were stained for 94 hours in 4ml of HCS Nuclear mask (Thermo Fisher UK) at concentration of 4 µL/mL in PBS at room temperature and with constant agitation. They were then bisected at the urethra, and the cut surface imaged on a glass-bottomed dish using the HREM microscope (all imaging parameters kept constant).

### 5.3 Resin testing

Multiple candidate resins were tested for setting time, compatibility with opacifying agents, final block hardness and quality of cuts. Resins used were Technovit 7100 (Heraeus Kulzer, Germany), Technovit 8100 (Heraeus Kulzer, Germany), Spurr resin (Polysciences Inc, USA), LR White (Sigma-Aldrich, USA) and Lowicryl HM20 (Polysciences Inc, USA). Composition of each resin is given in Supplementary Methods.

Hardness testing was conducted using either a Shore durometer D or Shore durometer A on set blocks. Cut quality was assessed by imaging a block with a mouse kidney embedded within it on the Optical HREM (Indigo Scientific, UK) and counting the number of slices which had areas of flaky resin or voids.

### 5.4 Spectroscopy

The transmission spectrum of OB was measured using an HG4000CG-UV-NIR (Ocean Optics) fiber-fed spectrometer. Samples of Technovit 8100 base sol. plus catalyst 1, with a low concentration (0.1 mg/mL) of OB, were measured in a PMMA semi-micro cuvette over a 4 mm path length, from 350 nm to 95 nm with a QTH10/M (Thorlabs) continuum lamp. The transmission spectrum (Figure 3C) has been compensated for the cuvette reflectivity and PMMA absorption.

### 5.5 Sub-surface fluorescence quantification

Standard 3D cell cultures were prepared as described (sup methods) and stained with HCS nuclear mask deep red or HSC nuclear mask. Samples were prepared for HREM as per Table 1, with Orasol Black 45X (Stort Chemicals Ltd, Bishops Stortford, UK) mixed with Technovit 8100 at the last stage of resin infiltration, (at concentrations of 2, 4, 8, 16 and 32 mg/mL) prior to positioning and setting the sample within the final block.

Blocks were imaged over the full sample depth on the Optical HREM system using slice thickness = 1.72 µm xy pixel size = 0.57 µm with gain = 9 and exposure = 1.0 s. Image stacks were down-sampled in xy to create isotropic voxels, background subtraction using rolling-ball algorithm with 50 pixelradius, and sliding parabola was used to remove autofluorescent background (ImageJ). Image stacks were resliced into xz stacks. ROI’s were manually drawn around isolated cells, fully enclosing all pixels with intensity above the background. The intensity profile of the ROI was calculated by averaging the intensity over the ROI columns (z-direction). This average signal was truncated at the maximum intensity and fit to a single exponential model (**Equation. 1**) in Matlab using a non-linear least squares approach, with start values of 0.5 for *I*_*0*_ *and τ*and limits of ± ∞ (other start values used made no difference to final fit parameters). The fit parameters, decay constant (*τ*) and initial intensity (*I*_*0*_), were used to quantify the change in subsurface fluorescence.

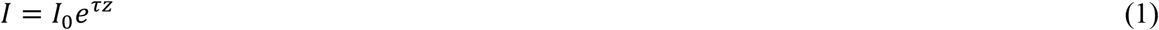

### 5.6 Image Post-processing PSF fitting and Deconvolution

HREM image stacks are inherently aligned and hence can be reconstructed as a volume without any z registration, alignment or overlap. Depending on the fluorophores used for labelling and the excitation emission filters used in the instrument, spectral unmixing (in the case of overlapping labelling spectra) or background subtraction to remove autofluorescence (in the case of shorter wavelength emission) may improve image appearance and aid in further segmentation (**Figure S2**).

For deconvolution the PSF parameters are estimated from the image stack as follows: The image stack was manually inspected for suitable small structures; these were cropped from the stack and a 3D median filtering with 1 pixel window size applied to reduce noise.^[28]^ Maximum intensity projections of the small structures in the x,y and z planes were used to estimate the mean FWHM in xy and z. A PSF was generated using a Gibson and Lanni model with parameters optimized based on minimizing the difference between measured and modelled FWHM (see Figure S2 and [11] for further details of PSF fitting). The PSF is dependent on many factors including but not limited to: the wavelength of the fluorophore, the concentration of opacifying agent, and pixel size (zoom level) of the microscope. Final PSF parameter values for each image stack with imaging parameters are given in Table S1. In the final stage the PSF was made asymmetric (due to physical sectioning discussed) by setting the lower half to zero and subtracting the background in the upper half (pixel intensities less than 0.0001 in 32 bit images were zeroed). This PSF was then used in a GPU accelerated RL deconvolution algorithm^[29]^ with border size of 1/4 of the image stack in each dimension respectively. Optimization of iterations was based on orthogonality of structures and contrast-to-noise ratio in the deconvolved final image stack.^[11]^

### 5.7 Murine tumor xenograft model

Eight-to ten-week-old, female, immune-compromised nu/nu nude mice (background CD1) were used (Charles River Laboratories). Cells from the FaDu cell line (a hypopharyngeal cancer cell line gifted by Dr Craig Murdoch (Sheffield University)) were cultured in complete medium (Dulbecco’s minimum essential medium Eagle with L-glutamine (DMEM) (Lonza) + 10% fetal bovine serum (Invitrogen)) in the ratio 1:10 (vol/vol) and incubated at 37 °C and 5% CO_2_. To prepare for injection, cells were washed with Dulbecco’s phosphate buffered saline and detached with trypsin-EDTA (7–8 min, 37 °C, 5% CO_2_) (Sigma). Cells were labelled with CMDiI (Thermofisher UK) a medium-term fluorescent cell-tracking dye that endures for approximately 3-6 cell divisions, and is transferred through cell division (but not cell-cell contact). Stain was dissolved from stock concentration (1mg/ml in Ethanol) in D-PBS to a working solution of 1 µM. Cells were incubated in the working solution for 5 minutes at 37°C, and then for 15 minutes at 4°C. Cells were then washed and re-suspended in PBS for injection. A 100 µl bolus of 1 × 10^6^ cells was injected subcutaneously into the left flank above the hind leg of each mouse (N=5), unstained cells were injected into the right flank also in a 100 µl bolus of 1 × 10^6^ cells. Tumor growth was measured daily with calipers every day after tumor became palpable, and were grown until total tumor volume was 1500mm^3^ or three weeks post-injection had elapsed.

For blood vessel staining, 200 µl Lectin (*Tomato*) bound to DyeLyte-649 (Vector UK) (1 mg/ml) was administered via tail vein injection and allowed to circulate for 10 minutes before perfusion fixation to allow sufficient binding to the vascular endothelium.^[3]^

### 5.8 Applications segmentation

For each application - tumor, kidney and brain - an example segmentation for biological structure of interests was carried out to demonstrate the quantification potential of MF-HREM data.

Glomeruli segmentation was performed in Vaa3D (v3.601) using the Gradient Vector Flow algorithm,^[30–32]^ with diffusion iteration of 5. This algorithm is a widely used extension to a traditional active contour segmentation technique, where the external energy term in the traditional active contour algorithm is replaced with the gradient vector flow field. Manual segmentation of the kidney cortex was then performed to remove structures not within the cortex (i.e. not glomeruli). Finally, a connected component analysis was performed in Amira v 19.2 with a threshold max size of 20000 µm^3^. No deconvolution was necessary (as measured by orthogonality ratio) owing to the size of the structures of interest (glomeruli) relative to the pixel size.

For the tumor, following deconvolution (parameters in Table S1), vascular segmentation was performed via the APP2 algorithm of Vaa3D (v3.601).^[30–33]^ Parameters were set as follows: threshold =-1(auto thresholding), CNN=3, GSBT was used and other parameters were used at their default values.

Brain microvasculature segmentation was performed after deconvolution, semi-manually in Amira-Avizo V 2019.2 using a 3D magic wand tool. White matter orientation analysis was performed using the OrientationJ plugin of Fiji^[34–36]^ with a Gaussian gradient and kernel sizes of 8 pixel or 4 pixel for Figures 7D and 7F respectively.

### 5.9 HREM imaging

All MF-HREM imaging was carried out using the Optical-HREM instrument (Indigo Scientific, UK) as shown in Figure 1. Samples were prepared into blocks as per Table 1. Blocks were then mounted into the instrument and initially sectioning without imaging until a flat block surface perpendicular to the optical axis was created. Once this surface was achieved, the desired slice thickness, x,y resolution, focus, total imaging depth, and exposure and gain for each wavelength were set. An air blower and vacuum were positioned to remove serial slices after sectioning. Imaging then proceeded in a fully automated manner until the total imaging depth had been achieved. For each application, imaging parameters are listed in Table S1.

## Supporting Information

Supporting Information is available from the Wiley Online Library or from the author.

## Acknowledgements

We would like to thank Craig Murdoch for the kind gift of the FaDu cell line.

## Table of Content text

### Multi-Fluorescence High-Resolution Episcopic Microscopy (MF-HREM) for Three-Dimensional Imaging of Adult Murine Organs

MF-HREM realizes the potential of optical HREM to image whole mouse organs with multiple fluorescence labels in 3D. This simple and robust serial-sectioning block-face technique is demonstrated in a range of biological contexts with proposed quantification pipelines.

N. A. Holroyd, E. Finnerty, C. Walsh*

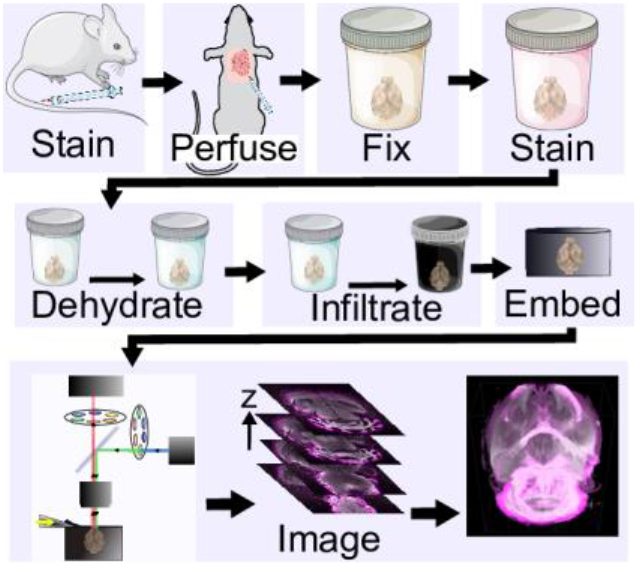

## References

1. N. Renier, Z. Wu, D. J. Simon, J. Yang, P. Ariel, M. Tessier-Lavigne, Cell 2014, 159, 896–910.

2. F. F. Voigt, D. Kirschenbaum, E. Platonova, S. Pagès, R. A. A. Campbell, R. Kastli, M. Schaettin, L. Egolf, A. van der Bourg, P. Bethge, K. Haenraets, N. Frézel, T. Topilko, P. Perin, D. Hillier, S. Hildebrand, A. Schueth, A. Roebroeck, B. Roska, E. T. Stoeckli, R. Pizzala, N. Renier, H. U. Zeilhofer, T. Karayannis, U. Ziegler, L. Batti, A. Holtmaat, C. Lüscher, A. Aguzzi, F. Helmchen, Nat. Methods 2019, 16, 1105.

3. A. d’Esposito, P. W. Sweeney, M. Ali, M. Saleh, R. Ramasawmy, T. A. Roberts, G. Agliardi, A. Desjardins, M. F. Lythgoe, R. B. Pedley, R. Shipley, S. Walker-Samuel, Nat. Biomed. Eng. 2018, 2, 773.

4. T. Liebmann, N. Renier, K. Bettayeb, P. Greengard, M. Tessier-Lavigne, M. Flajolet, Cell Rep. 2016, 16, 1138.

5. J. A. Gleave, J. P. Lerch, R. M. Henkelman, B. J. Nieman, PLoS One 2013, 8, e72039.

6. K. Tainaka, T. C. Murakami, E. A. Susaki, C. Shimizu, R. Saito, K. Takahashi, A. Hayashi-Takagi, H. Sekiya, Y. Arima, S. Nojima, Cell Rep. 2018, 24, 2196.

7. J. H. Kim, M. J. Jang, J. Choi, E. Lee, K. D. Song, J. Cho, K. T. Kim, H. J. Cha, W. Sun, Sci. Rep. 2018, 8, 1.

8. T. Yu, Y. Qi, H. Gong, Q. Luo, D. Zhu, J. Biophotonics 2018, 11, e201700187.

9. D. G. C. Hildebrand, M. Cicconet, R. M. Torres, W. Choi, T. M. Quan, J. Moon, A. W. Wetzel, A. S. Champion, B. J. Graham, O. Randlett, Nature 2017, 545, 345.

10. G. Krishnamurthi, C. Y. Wang, G. Steyer, D. L. Wilson, Opt. Express 2010, 18, 22324.

11. C. Walsh, N. Holroyd, R. Shipley, S. Walker-Samuel, in Ann. Conf. MUIA 2020, 235.

12. S. P. Amato, F. Pan, J. Schwartz, T. M. Ragan, Front. Neuroanat. 2016, 10, 1–11.

13. K. Seiriki, A. Kasai, T. Nakazawa, M. Niu, Y. Naka, M. Tanuma, H. Igarashi, K. Yamaura, A. Hayata-Takano, Y. Ago, H. Hashimoto, Nat. Protoc. 2019, 14, 1509.

14. H. Gong, D. Xu, J. Yuan, X. Li, C. Guo, J. Peng, Y. Li, L. A. Schwarz, A. Li, B. Hu, B. Xiong, Q. Sun, Y. Zhang, J. Liu, Q. Zhong, T. Xu, S. Zeng, Q. Luo, Nat. Commun. 2016, 7, 1.

15. L. Abdeladim, K. S. Matho, S. Clavreul, P. Mahou, J.-M. Sintes, I. Solinas, Xavier Arganda-Carreras, S. G. Turney, J. W. Lichtman, A. Chessel, K. Bemelmans, Alexis-Pierre Loulier, W. Supatto, J. Livet, E. Beaurepaire, Nat Commun. 2019, 10, 1662.

16. S. H. Geyer, B. Maurer-Gesek, L. F. Reissig, W. J. Weninger, J. Vis. Exp. 2017, 125, e56071.

17. W. J. Weninger, S. H. Geyer, A. Martineau, A. Galli, D. J. Adams, DMM 2014, 7, 1143.

18. G. Pieles, S. H. Geyer, D. Szumska, J. Schneider, S. Neubauer, K. Clarke, K. Dorfmeister, A. Franklyn, S. D. Brown, S. Bhattacharya, W. J. Weninger, J. Anat. 2007, 211(1), 132.

19. W. J. Weninger, S. H. Geyer, T. J. Mohun, D. Rasskin-Gutman, T. Matsui, I. Ribeiro, L. da F. Costa, J. C. Izpisua-Belmonte, G. B. Muller, Anat. Embryol. (Berl). 2006, 211, 213.

20. T. Mohun, D. J. Adams, R. Baldock, S. Bhattacharya, A. J. Copp, M. Hemberger, C. Houart, M. E. Hurles, E. Robertson, J. C. Smith, T. Weaver, W. J. Weninger, DMM, 2013, 6, 562.

21. J. Rosenthal, V. Mangal, D. Walker, M. Bennett, T. J. Mohun, C. W. Lo, Birth Defects Res. C Embryo Today 2004, 72, 213.

22. T. J. Mohun, W. J. Weninger, Curr. Opin. Genet. Dev. 2011, 21, 573.

23. S. H. Geyer, M. M. Nöhammer, I. E. Tinhofer, W. J. Weninger, J. Anat. 2013, 223, 603.

24. H. Matsui, S. Y. Ho, T. J. Mohun, H. M. Gardiner, Ultrasound Obstet. Gynecol. 2015, 45, 492.

25. W. J. Weninger, T. J. Mohun, Methods Mol. Biol. 2007, 411, 35.

26. T. J. Mohun, W. J. Weninger, Cold Spring Harb. Protoc. 2012, 7, 678.

27. P. Workman, E. O. Aboagye, F. Balkwill, A. Balmain, G. Bruder, D. J. Chaplin, J. A. Double, J. Everitt, D. A. H. Farningham, M. J. Glennie, L. R. Kelland, V. Robinson, I. J. Stratford, G. M. Tozer, S. Watson, S. R. Wedge, S. A. Eccles, Br. J. Cancer 2010, 102, 1555.

28. J. Schindelin, I. Arganda-Carreras, E. Frise, V. Kaynig, M. Longair, T. Pietzsch, S. Preibisch, C. Rueden, S. Saalfeld, B. Schmid, J.-Y. Tinevez, D. J. White, V. Hartenstein, K. Eliceiri, P. Tomancak, A. Cardona, Nat. Methods 2012, 9, 676–682.

29. C. Rueden, C. Dietz, M. Horn, J. Schindelin, B. Northan, M. Berthold, K. Eliceiri, ImageJ Ops [software] 2016.

30. H. Peng, A. Bria, Z. Zhou, G. Iannello, F. Long, Nat. Protoc. 2014, 9, 193.

31. H. Peng, Z. Ruan, F. Long, J. H. Simpson, E. W. Myers, Nat. Biotechnol. 2010, 28, 348.

32. H. Peng, J. Tang, H. Xiao, A. Bria, J. Zhou, V. Butler, Z. Zhou, P. T. Gonzalez-Bellido, S. W. Oh, J. Chen, A. Mitra, R. W. Tsien, H. Zeng, G. A. Ascoli, G. Iannello, M. Hawrylycz, E. Myers, F. Long, Nat. Commun. 2014, 5, 4342.

33. H. Xiao, H. Peng, Bioinformatics 2013, 29, 1448–1454.

34. Z. Püspöki, M. Storath, D. Sage, M. Unser, Focus on Bio-Image Informatics, Springer, 2016, 69.

35. R. Rezakhaniha, A. Agianniotis, J. T. C. Schrauwen, A. Griffa, D. Sage, C. V. C. vd Bouten, F. N. Van De Vosse, M. Unser, N. Stergiopulos, Biomech. Model. Mechanobiol. 2012, 11, 461.

36. E. Fonck, G. G. Feigl, J. Fasel, D. Sage, M. Unser, D. A. Rüfenacht, N. Stergiopulos, Stroke 2009, 40, 2552.

37. D. J. Jafree, D. Moulding, M. Kolatsi-Joannou, N. P. Tejedor, K. L. Price, N. J. Milmoe, C. L. Walsh, R. M. Correra, P. J. D. Winyard, P. C. Harris, C. Ruhrberg, S. Walker-Samuel, P. R. Riley, A. S. Woolf, P. J. Scambler, D. A. Long, Elife 2019, 8, e48183.

38. W. Li, R. N. Germain, M. Y. Gerner, Proc. Natl. Acad. Sci. 2017, 114, e7321.

39. E. Chee Tak Yeung, Plant Microtechniques and Protocols, Springer, 2015.

40. Z. Yang, B. Hu, Y. Zhang, Q. Luo, H. Gong, PLoS One 2013, 8, 4.

41. Y. Gang, X. Liu, X. Wang, Q. Zhang, H. Zhou, R. Chen, L. Liu, Y. Jia, F. Yin, G. Rao, J. Chen, S. Zeng, Biomed. Opt. Express 2017, 8, 3583.

42. A. J. Ewald, H. Mcbride, M. Reddington, S. E. Fraser, R. Kerschmann, Dev. Dyn. 2002, 225, 369.

43. W. Wallace, L. H. Schaefer, J. R. Swedlow, Biotechniques 2001, 31, 1076.

44. F. Soulez, L. Denis, Y. Tourneur, É. Thiébaut, 9th IEEE International Symposium on Biomedical Imaging (ISBI) 2012, 1735.

45. D. Sage, L. Donati, F. Soulez, D. Fortun, G. Schmit, A. Seitz, R. Guiet, C. Vonesch, M. Unser, Methods 2017, 115, 28.

46. J. B. de Monvel, E. Scarfone, S. Le Calvez, M. Ulfendahl, Biophys. J. 2003, 85, 3991.

47. E. J. Baldelomar, J. R. Charlton, S. C. Beeman, B. D. Hann, L. Cullen-McEwen, V. M. Pearl, J. F. Bertram, T. Wu, M. Zhang, K. M. Bennett, Kidney Int. 2016, 89, 498.

48. A. Klingberg, A. Hasenberg, I. Ludwig-Portugall, A. Medyukhina, L. Männ, A. Brenzel, D. R. Engel, M. T. Figge, C. Kurts, M. Gunzer, J. Am. Soc. Nephrol. 2017, 28, 452.

49. D. Hanahan, R. A. Weinberg, Cell 2011, 144, 646.

50. P. Friedl, K. Wolf, Nat. Rev. Cancer 2003, 3, 362.

51. T. Magdeldin, V. López-Dávila, J. Pape, G. W. W. Cameron, M. Emberton, M. Loizidou, U. Cheema, Sci. Rep. 2017, 7, 1.

52. X. Zhang, X. Yin, J. Zhang, A. Li, H. Gong, Q. Luo, H. Zhang, Z. Gao, H. Jiang, Natl. Sci. Rev. 2019, 6, 1223.

53. J. Steinman, M. M. Koletar, B. Stefanovic, J. G. Sled, PLoS One 2017, 12(10), e0186676.

54. M. I. Todorov, J. C. Paetzold, O. Schoppe, G. Tetteh, V. Efremov, K. Völgyi, M. Düring, M. Dichgans, M. Piraud, B. Menze, A. Ertürk, (preprint) bioRxiv, 613257, submitted: April, 2019.

55. M. D. Budde, J. A. Frank, Neuroimage 2012, 63, 1.

56. R. Cai, C. Pan, A. Ghasemigharagoz, M. I. Todorov, B. Förstera, S. Zhao, H. S. Bhatia, A. Parra-Damas, L. Mrowka, D. Theodorou, M. Rempfler, A. L. R. Xavier, B. T. Kress, C. Benakis, H. Steinke, S. Liebscher, I. Bechmann, A. Liesz, B. Menze, M. Kerschensteiner, M. Nedergaard, A. Ertürk, Nat. Neurosci. 2019, 22, 317.

57. J. Kohl, J. Ng, S. Cachero, E. Ciabatti, M.-J. Dolan, B. Sutcliffe, A. Tozer, S. Ruehle, D. Krueger, S. Frechter, T. Branco, M. Tripodi, G. S. X. E. Jefferis, Proc. Natl. Acad. Sci. 2014, 111, e3805.

58. Y. Qi, T. Yu, J. Xu, P. Wan, Y. Ma, J. Zhu, Y. Li, H. Gong, Q. Luo, D. Zhu, Sci. Adv. 2019, 5, eaau8355.

59. J. Guo, C. Artur, J. L. Eriksen, D. Mayerich, Sci. Rep. 2019, 9, 14578.

60. D. K. Samuylov, P. Purwar, G. Szekely, G. Paul, IEEE Trans. Image Process. 2019, 28, 3688.

61. J. E. D. Zamboni, V. H. Casco, J. Imaging 2017, 3.

62. R. J. Vigouroux, M. Belle, A. Chédotal, Mol. Brain 2017, 10, 1.

