## Supplementary Methods for "Multi-Fluorescence High-Resolution Episcopic Microscopy (MF-HREM) for Three-Dimensional Imaging of Adult Murine Organs"

### SI 1. Supplementary methods

#### SI 1.1 3D cell culture for standardised samples

Compressed type I collagen hydrogels (RAFT UK) were used as standardized samples for testing stain compatibility with tissue processing and for quantification of subsurface fluorescence. Samples were prepared in a 24-well plate using the protocol described by the manufacturer, with the addition of SW1222 colorectal cancer cells at 100,000 cells/mL (25). Samples were fixed with 4% PFA for 20 mins. For subsurface fluorescence experiments, post-fixation, cell nuclei were stained with the addition of HCS nuclear mask deep red 2 $\mu$L/mL in PBS incubated for 30 mins. Samples were then processed as described in Table 1.

#### SI 1.2 Resin testing

- Technovit 7100 is a 2-hydroxyethyl methacrylate-based plastic resin and was prepared using 1 g Technovit 7100 hardener 1 dissolved in 100 mL Technovit 7100 resin. Technovit 7100 hardener 2 was used the catalyze the polymerisation reaction, added in a ratio of 1:15 Hardener 2 to resin.
- Technovit 8100 had a similar composition: 0.5 g of Technovit 8100 Hardener 1 dissolved in 100 mL Technovit 8100 resin. The catalyst, Technovit 8100 Hardener 2, was added in a ratio of 1:30 catalyst to resin.
- Spurr is an epoxy resin, and was prepared using 4.1 g ERL, 1.43 g diglycidyl ether of polypropylene glycol, 5.9 g nonenylsuccinic anhydride and 0.1 g dimethylaminoethanol accelerator.
- LR White is an acrylic resin and was prepared using 2 g benzoyl peroxide accelerator per 100 mL resin.
- Lowicryl HM20, a methacrylate resin, was prepared with 0.6% (w/w) benzoyl peroxide accelerator.

#### SI 1.3 Stain Penetration

Freeze-Thaw: Kidneys were dehydrated through a methanol in $dH_2O$ series: 20%, 40%, 60%, 80%, 100% for 1 hr in each (7ml per kidney). Kidneys were freeze-thawed 3 times for 20 mins each time at -80 °C. Kidneys were then rehydrated through methanol series 80%,60%,40%,20% ,0% (1 hr each).

iDISCO: Kidneys were washed in PTX.2 1hr two times at room temp. Kidneys were incubated overnight at 37°C in a solution containing 1xPBS, 0.2% Triton-X (Sigma UK), 20% DMSO (Sigma UK). Kidneys were then incubated overnight at 37°C in a solution of 1xPBS, 0.1% Tween-20 (Sigma UK), 0.1% Triton-X, 0.1% Deoxycholate (Sigma UK), + 0.1% NP40 (Sigma UK), 20% DMSO.

P[K]: Tris Buffer containing 1.21 g Tris (Sigma UK), 0.147 g $CaCl_2.H_2O$, (Sigma UK), 65 ml dH2O, 30 ml glycerol (Sigma UK) was made and 40 µg/mL proteinase [K] (Sigma UK) was added. The sample was incubated at room temp for 10 mins, with constant agitation before being washed in PBS x3 for 10 mins each.

Saponin: A solution containing 2 g of gelatin (VWR) in 1 L PBS was made and filtered immediately. After allowing the solution to chill, 5 mL of Triton X-100, 0.1 g of sodium azide (Sigma UK) and 10 mg/mL saponin (Sigma UK) was added to the solution. Kidneys were incubated for (72 hrs) in 3 ml of solution (PBS for control kidneys) at room temp. with constant agitation.

### SI 1.4 Imaging parameters

| Sample | FaDU Tumour | WT Kidney | Brain 1 | Brain 2 |
|---|---|---|---|---|
| Structures stained | 1. Injected cells<br>2. vasculature | 1. Vasculature<br>2. cell membrane<br>3. Nucleus, cytoplasm, Cytosol | 1. Vasculature<br>2. White matter | 1. White matter<br>2. Cell nuclei |
| fluorephores excitation/emission (nm) | 1. CMDiI (553⁄570)<br>2. Dylight649 (655/670) | 1. Dylight649 (655/670)<br>2. CMDiI (553⁄570)<br>3. HCS cell mask blue (346/422) | 1. Dylight649 (655/670)<br>2. CMDiI (553⁄570) | 1. CMDiI (553⁄570)<br>2. HCS cell mask deep red (638⁄686) |
| Imaging time | 12 hrs | 12 hrs | 12 hrs | 19hrs |
| No. of sections | 1628 | 1599 | 1524/314 | 1954 |
| Pixel size (x,y,z) (µm) | 2.75, 2.75, 2.58 | 2.17, 2.17, 2.58 | 2.75,2.75,2.58<br>zoom region - 0.57,0.57,1.72 | 4.5,4.5,2.58 |
| Conc. Orasol Black (mg/mL) | 1 | 4 | 4 | 1 |
| PSF model | Gibson&Lanni | Na | Gibson& Lanni | Na |
| PSF parameters | Parameter \| 1 \| 2<br>$RI_{sample}$ \| 1.5 \| 1.5<br>$RI_{immersion}$ \| 1 \| 1<br>NA \| 0.1 \| 0.165<br>Offset \| 0 \| 0<br>working dist. \| 150 \| 150<br>Wavelength \| 2800 \| 2175 | Na | $RI_{sample}$ = 1.5<br>$RI_{immersion}$ = 1.0<br>NA = 0.25<br>Offset = 0µm<br>working dist.=150µm<br>Wavelength= 1200nm | Na |
| Segmentation/quantification technique | 1. Otsu Threshold<br>2. APP2 | 1.Gradient vector flow<br>2. Na<br>3. Na | 1.3D magic wand tool.<br>2. OrientationJ | 1. OrientationJ<br>2. Na |
| Deconvolution iteration no. | 1. 30<br>2. 40 | Na | 1. 20<br>2. Na | Na |

**Table S1**. Detailing the imaging parameters for all samples

### S 2. Supplementary Results

| Stain | Fluorescence retained in ethanol | Fluorescence retained in acetone | Manufacture/Supplier and catalogue number |
|---|---|---|---|
| Eosin B | Yes | Yes | Sigma 45260 |
| Eosin Y | (Poor solubility) | NT | Sigma 230251 |
| Acridine Orange | Yes | Yes | Sigma A6014 |
| Actin Green™ 488 Ready Probes | No | NT | Thermo Fisher R37110 |
| NucRed™ Live 647 ReadyProbes | No | NT | Thermo Fisher R37106 |
| CellMask™ Orange Plasma membrane Stain | No | NT | Thermo Fisher C10045 |
| HCS CellMask™ Red Stain | Yes | Yes | Thermo Fisher H32712 |
| HCS NuclearMask™ Deep Red Stain | Yes | Yes | Thermo Fisher H10294 |
| DAPI | Yes | No | Sigma Aldrich D9542 |
| Invitrogen™ Lectin GS-II From *Griffonia simplicifolia*, Alexa Fluor™ 647 Conjugate | Yes | Yes | Invitrogen™ L32451 |
| DyLight 649 labeled Lycopersicon Esculentum (Tomato) Lectin (LEL, TL) | Yes | Yes | Vector DL-1178-1 |
| Anti-Neurofilament heavy polypeptide antibody | Yes | No | Abcam ab4680 |
| CellTracker™ CM-DiI Dye | Yes | Yes | Invitrogen C7001 |
| Propidium iodide | yes | NT | Invitrogen P1304MP |
| SP-DiO$C_{18}$(3) (3,3'-Dioctadecyl-5,5'-Di(4-Sulfophenyl)Oxacarbocyanine, Sodium Salt) | Poor solubility | Poor solubility | Invitrogen D7778 |
| Wheat Germ Agglutinin, Alexa Fluor™ 647 Conjugate | Yes | Yes | Invitrogen W32466 |
| Green fluorescent protein (GFP) | No | No | NA |

**Table S2**. Stain compatibility with various dehydrants

### S 2.2 Resin testing

| Resin | Dehydratant | Setting temp | Oxygen | Time to set w/wo opacifying agent | Slice cut quality | Observations |
|---|---|---|---|---|---|---|
| **Technovit 7100** | Ethanol | RT | Room | 10(mins)/1(hr) | + | |
| **Technovit 8100** | Acetone | 4°C | Vacuum/mineral Oil | 10/30 (mins) | ++ | |
| **Lowicryl HM20** | Ethanol | RT°C | Room | NA/NA | NA | Did not set after 120hrs |
| **Spurr** | Ethanol | 55°C | Room | 1.9/2.2(hrs) | - | All blocks shattered during cut testing |
| **LR White** | Ethanol | 55°C | Room | 20/23 (hrs) | NA | Large amount of resin expansion upon setting |

**Table S3.** Resin testing results

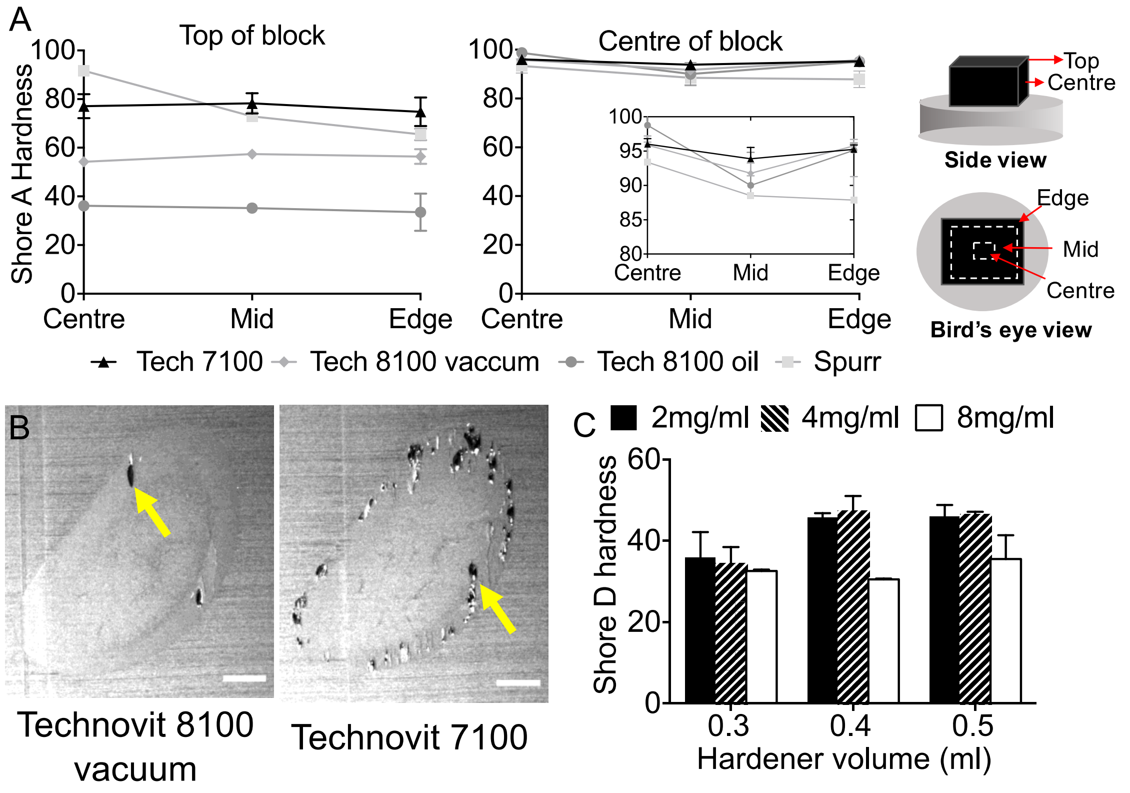


**Figure S1.** Characterisation of potential resins and optimisation of embedding procedure. A) Hardness measurements for 3 candidate resins (Technovit 7100, Technovit 8100 and Spurr with Technovit 8100 under two different oxygen exclusion conditions: oil or vacuum. Hardness measurements were made using a Shore Duromenter A at two axial positions: top surface and block centre, and at 3 lateral positions for each axial: centre, mid and edge (as indicated in the diagram. Measurements were made in triplicate on two independent samples for each case (mean and standard deviation shown). For all resins shown blocks were harder at the centre of the block than the top of the block (axial position). There was a gradient of decreasing hardness for Spurr resin from the centre to the edge. Technovit 8100 set under oil had the lowest hardness at the top of the block but in the centre all resins had similar measured hardness. B) Automated cut quality was assessed using the HREM, for Technovit 7100 and 8100 the proportion of slices with flakey resin or voids was used as a metric for cut quality. Representative images for Technovit 8100 and Technovit 7100, showing the flaky resin/voids (yellow arrows) (scale bar =1mm). C) Hardness measurements for Technovit 8100 set under vacuum with three different amounts of secondary catalyst (0.3, 0.4 or 0.5 mL per 15 mL of infiltration soln.) and Orasol Black (2,4,8 mg/mL). Hardness was measured on the top of the block (axially) and in the centre of the block (laterally) with 2 replicate measures on two independent samples in each case. Results show a significant increase in hardness with increasing secondary catalyst and decreasing concentration of Orasol Black (Two-way anova $p=0.0059$ for secondary catalyst $p=0.0011$ for opacifying agent concentration.)

### S 2.3 Image processing

Depending on the fluorophores used for labelling and the excitation emission filters used in the instrument, spectral unmixing (in the case of overlapping labelling spectra) or background subtraction to remove autofluorescence (in the case of shorter wavelength emission) may improve signal to noise ratio, providing clearer imaging as potentially aiding in further segmentation.

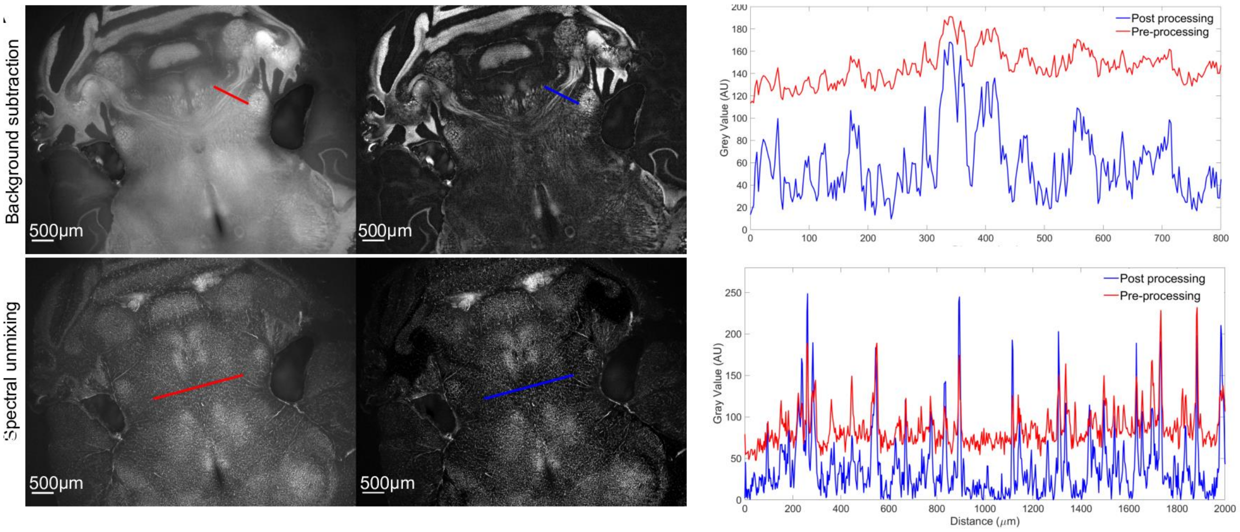


**Figure S2.** Showing the increase in signal-to-noise ratio achieved by pre-processing, using a rolling-ball algortihm to remove autofluorescence (upper row) and spectral unmixing to remove cross talk in multi-fluorescent image stacks (lower row)
